# CIRCE: a scalable Python package to predict cis-regulatory DNA interactions from single-cell chromatin accessibility data

**DOI:** 10.1101/2025.09.23.678054

**Authors:** Rémi Trimbour, Julio Saez-Rodriguez, Laura Cantini

**Affiliations:** Institut Pasteur, Université Paris Cité, CNRS UMR 3738, Machine Learning for Integrative Genomics Group, F-75015 Paris, France; Heidelberg University, Faculty of Medicine, and Heidelberg University Hospital, Institute for Computational Biomedicine, Heidelberg, Germany; European Molecular Biology Laboratory, European Bioinformatics Institute (EMBL-EBI),Hinxton, Cambridgeshire, U.K.

## Abstract

Chromatin 3D folding creates numerous DNA interactions, participating in gene expression regulation. Single-cell chromatin-accessibility assays now profile hundreds of thousands of cells, challenging existing methods for mapping cis-regulatory interactions.

We present CIRCE, a fast and scalable Python package to predict cis-regulatory DNA interactions from single-cell chromatin accessibility data. CIRCE re-implements the Cicero workflow to analyse single-cell atlases, cutting runtime and memory use by several orders of magnitude. We also provide new options to compute metacells, grouping similar cells to reduce data sparsity.

We benchmarked CIRCE against Cicero on two datasets of different sizes and demonstrated the improvement from CIRCE’s metacells’ strategy with promoter capture Hi-C data. We also evaluated how DNA interaction predictions are impacted by different pre-processing. We observed a negative impact of Cicero’s count normalization, and the best performance was obtained with the single-cell count matrix directly. Finally, we demonstrated the scalability of CIRCE by processing a dataset of more than 700000 cells and 1 million DNA regions in less than an hour.

CIRCE should greatly facilitate the prediction of DNA region interactions for scverse and Python users, while providing new and up-to-date pre-processing insights.

**Availability and reproducibility:** CIRCE is released as an open-source software under the AGPL-3.0 license. The package source code is available on GitHub at https://github.com/cantinilab/CIRCE, and its documentation is accessible at https://circe.readthedocs.io.

The code to reproduce the presented results is available as a Snakemake pipeline at https://github.com/cantinilab/circe_reproducibility.

## 1. Introduction

Cis-regulatory elements can regulate genes located hundreds of thousands of base pairs away through DNA folding, which brings distal regulatory elements into close spatial arrangement with the gene promoter region to facilitate interactions (Dekker *et al*., 2002; Lieberman-Aiden *et al*., 2009). The regulation of gene expression results from complex and dynamic interactions between all these regulatory regions.

Single-cell Assay for Transposase-Accessible Chromatin using sequencing (scATAC-seq) is a powerful technique for studying chromatin accessibility in individual cells. The chromatin accessibility profiles measured allow to understand the role of specific DNA elements, such as enhancers, to define cell state heterogeneity and regulate gene expression (Badia-i-Mompel *et al*., 2023).

While single-cell ATAC-seq provides insights into chromatin accessibility and potential interactions with transcription factors and the transcriptional machinery, identifying regulatory and interacting regions from chromatin accessibility remains challenging, as it does not inform about DNA conformation.

Several methods took interest in this challenge and proposed to infer gene-enhancer connections from single-cell ATAC-seq data alone (Stuart *et al*., 2021; Granja *et al*., 2021; Pliner *et al*., 2018), single-cell multi-omics data (Zhang *et al*., 2022; Bravo González-Blas *et al*., 2023; Trimbour *et al*., 2024; Sakaue *et al*., 2024), or combining ATAC with other input data types, such as Hi-C data or DNA sequences (Fulco *et al*., 2019; Grover *et al*., 2024; Sheth *et al*., 2024). Among them, Cicero, an algorithm developed for analysing single-cell ATAC-seq data (Pliner *et al*., 2018), has been widely adopted to uncover cis-regulatory DNA interactions and gene regulatory mechanisms (Kamimoto *et al*., 2023; Trimbour *et al*., 2024). It aims to construct a global map of cis-regulatory interactions, considering the distance between regions and the technical effects in measurements.

Cicero was developed when single-cell datasets were relatively small and optimal preprocessing strategies had not been well established. Cicero shows limited resource usage performances when applied to very large high-resolution datasets produced nowadays (e.g., hundreds of thousands of cells) (Cusanovich *et al*., 2018; Yao *et al*., 2021; Zhang *et al*., 2021). Its single-cell preprocessing also relies by default on a clustering of cells from UMAP/t-SNE spaces, which do not accurately preserve distances between observations (Chari and Pachter, 2023; Kobak and Linderman, 2021). Furthermore, Cicero is only available as an R package, and integrating it in a Python workflow, in particular in the broadly used scverse ecosystem, requires hybrid environments and additional effort. For example, both gene regulatory network inference methods, HuMMuS (Trimbour et al., 2024) and CellOracle (Kamimoto et al., 2023), are currently based on Cicero outputs and require users to export them to follow the next Python workflow steps.

To overcome these limitations, we introduce CIRCE, a fast and scalable Python package to analyse single-cell ATAC datasets and atlases. We update the metacells computation strategy and preprocessing following recent literature guidelines, benchmarking each strategy’s performance. CIRCE implements the algorithm proposed in (Pliner *et al*., 2018) and initially available in the R package Cicero cole-trapnell-lab.github.io/cicero-release. CIRCE runs ∼150 times faster, allowing CPU parallelisation, and uses significantly less memory. CIRCE allows users to integrate cis-regulatory network computation into scverse workflows (Virshup *et al*., 2023) and considerably decreases memory usage for all dataset sizes.

We also provide some insights into the impact of input data preparation. Aggregating highly similar cells into “metacells” can mitigate the sparsity and sampling noise inherent to single-cell profiles such as scRNA-seq, while reducing dataset size (Baran *et al*., 2019; Bilous *et al*., 2024). Due to its extreme sparsity, scATAC is often analysed after metacells computation (Persad *et al*., 2023; Pliner *et al*., 2018). We compared cis-regulatory interaction predictions from single-cell versus metacell inputs, as well as binarized versus un-binarized counts. Using promoter capture Hi-C data, we observed higher predictions with CIRCE metacells than Cicero metacells. However, we show that using directly single-cell input data led to even better predictions despite their higher sparsity.

While metacells are particularly interesting for condensing information from large datasets and reducing computational requirements, there seems to be a trade-off between resource usage and small performance improvement. CIRCE offers both an improved strategy for metacell computation and the capacity to use single-cell as input up to atlas-sized datasets, by greatly improving processing speed and memory usage.

## 2. Implementation

### 2.1. Workflow and parameters

The methodologies of CIRCE and Cicero differ in both the preprocessing of the scATAC-seq data and the strategy for grouping highly similar cells in metacells before computing the co-accessibility scores. In its last implementation, Cicero proposes by default to compute latent semantic indexing (LSI) reduction from the binarized input data, projecting the cells into a lower-dimensional space of DNA region-topics, summarizing the main chromatin patterns. Cicero then computes a second dimensionality reduction on this space (UMAP or t-SNE), from which it calculates the nearest neighbours. It has now been demonstrated that t-SNE or UMAP spaces, especially when keeping very few dimensions, do not conserve distance between observations and thus are not adapted to compute nearest neighbours (Chari and Pachter, 2023). In CIRCE, we propose by default to group the cells directly on a reduction space generated with LSI, while still providing the option to compute a UMAP reduction and re-use its clustering strategy.

CIRCE relies on the algorithm proposed in (Pliner *et al*., 2018) to compute co-accessibility, which uses Graphical Lasso to impose a distance penalization on the covariance of DNA regions (see Figure 3.1.a). To reduce computational complexity, these calculations are performed within a sliding window, where the window size corresponds to the maximum distance considered for cis-interactions. This limit is by default 500kb for human and mouse cis-regulatory interactions in both Cicero and CIRCE, a distance also used as a window limit around promoters to find distal enhancers in several other works (Forrest *et al*., 2014; Stuart *et al*., 2021). Distance penalties are computed between every pair of DNA regions according to the following formula *ρ* = (*1* − ^−^) ×, with the distance between two regions.

The scaling exponent of the power-law function estimates the global decay in contact frequency. It has been estimated at 0.75 for the “tension globule” polymer model in human and mouse (Sanborn *et al*., 2015), and a value of 0.85 has been proposed for Drosophila (Sexton *et al*., 2012). The parameter α, proportional to the penalty, enforces the sparsity of the final result. It is estimated by selecting random windows and calculating the lowest value for each of them, such that less than 5% of the long-range pair of DNA regions (usually >250kb distant DNA regions) and 80% of all pairs in the window are non-zero entries. To simplify the choice of the species-specific parameters in CIRCE (, window size, distance of long and short-range interactions), users can also specify the organism corresponding to their data to select automatically the default literature-based values.

Finally, cis-co-accessibility networks (CCANs) are defined as described in (Pliner *et al*., 2018), using the Louvain clustering method (Blondel *et al*., 2008) and the same default parameters.

### 2.2. Implementation details

The most popular Graphical Lasso model (the core of CICERO) in Python is hosted in the scikit-learn (Pedregosa *et al*., 2011) package, but it does not allow a pairwise matrix penalty. Consequently, we use the skggm package (Laska and Narayan, 2017), implementing the QUIC algorithm (Hsieh *et al*., 2014) in a scikit-learn compatible package and offering more freedom on model penalization. CCANs identification is based on the Louvain clustering method implemented in the NetworkX package (Hagberg *et al*., 2008).

CIRCE works directly on AnnData objects (Virshup *et al*., 2024), making it fully compatible with any scVerse package (Virshup *et al*., 2023). All results are stored as a sparse matrix for memory usage optimization, directly in the ‘.obsp’ slot of your input AnnData object, and can easily be extracted as a readable table.

Additionally, we provide a tutorial to extend the co-accessible DNA region networks of CIRCE by building prior gene regulatory networks for CellOracle. This tutorial is accessible at https://circe.readthedocs.io/en/latest/examples/4_circe_celloracle_tutorial.html.

## 3. Performances and comparison with Cicero

We compared CIRCE and Cicero with and without generating metacells, on a BMMC dataset (1,771 cells and 110,235 peaks), and a BPMC dataset (9,631 cells and 215,676 peaks). We demonstrated that both implementations were returning near-identical results from the same input, then explored the impact of respective preprocessing strategies on inferring enhancer-promoter interactions.

**Figure 1.**
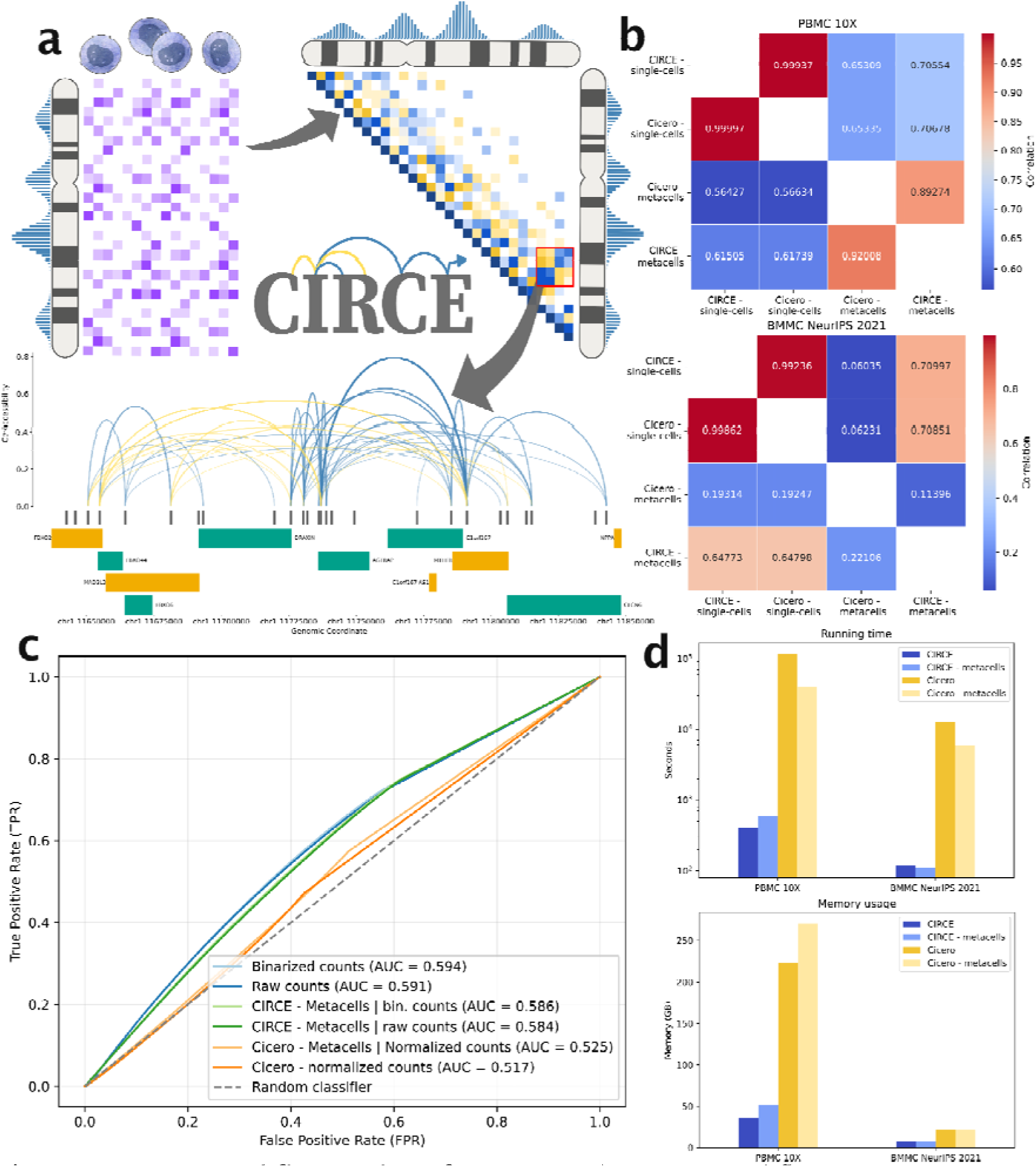
CIRCE workflow and performances. **a)** CIRCE workflow overview. From scATAC-seq data, co-accessibility scores are defined as regularized covariance values between DNA regions, using a graphical lasso model and pairwise distance penalties. Cis-coaccessible networks (CCANs) can then be extracted with the Louvain community detection method, as modules of DNA regions with high absolute co-accessibility scores. Co-accessibility scores in a CCAN or a DNA window of interest can then be visualised with different graphical options. **b)** Correlation values between the networks obtained from CIRCE, Cicero, and metacells computation on both the BMMC and the PBMC datasets. The upper triangle of the heatmap contains the Spearman correlation, while the lower triangle contains the Pearson correlation. Colour gradient illustrates the correlation values. **c)** ROC curves on the recovering DNA region-promoter interactions obtained from the PC-HiC dataset. Different preprocessed inputs are evaluated for each method: CIRCE from raw single-cell counts, binarized single-cell counts, and metacell counts from LSI dimensionality reduction space on both single-cell count matrices, and Cicero on the corrected count matrices and the metacell count matrices obtained from the binarized corrected counts. **d)** Running-time and memory-usage of CIRCE and Cicero when running on the single-cell or computing metacells.

From the same single-cell ATAC-seq input data, we observed nearly identical results, with respective Spearman and Pearson correlations of 0.9923 and 0.9986 in the BMMC datasets, and 0.9993 and 0.9999 on the PBMC dataset (Figure 3.1b). The small difference can be explained by the stochasticity in the algorithm that selects random DNA windows to estimate the parameter α (see Implementation section).

However, we observed substantial differences when using the respective metacell strategies of CIRCE and Cicero. First, CIRCE’s metacells were still relatively highly correlated with single-cell inferred networks, with respective Spearman and Pearson correlations of 0.71 and 0.65 on the BMMC dataset, and 0.71 and 0.62 on the PBMC datasets. In contrast, the correlations with Cicero’s metacells were lower (Figure 3.1b), with particularly high variability between the datasets. On the BMMC dataset, the correlations with the single-cell data run were especially low (i.e., Spearman and Pearson correlations respectively 0.06 and 0.19), while the decrease was less marked on the PBMC dataset (i.e., Spearman and Pearson correlations of 0.65 and 0.57).

After observing such differences in the predicted DNA region interactions, we decided to evaluate the networks generated to identify the best strategy. Using a Promoter Capture Hi-C dataset (Javierre *et al*., 2016), we measured how well each method was recovering enhancer-promoter interactions (Figure 3.1c). Surprisingly, the networks with the highest AUCs were obtained from the binarized single-cell count matrix and the un-binarized single-cell count matrix (respectively 0.594 and 0.591). In comparison, the predictions from the default corrected counts with Cicero’s function make_atac_cds had an AUC of 0.517. We tested the default metacell strategy of CIRCE on both raw and binarized counts, both leading to a small decrease in the AUC value, with both the binarized and un-binarized metacell counts performing almost identically (0.586 compared to 0.584). The prediction from Cicero’s metacells gave an AUC of 0.525, or a small improvement compared to the predictions from Cicero’s processing of single-cell input. Overall, we observed better enhancer-promoter interaction predictions when using the single-cell matrix directly, and without Cicero’s count correction. This result suggests that Cicero’s count correction is not the best preprocessing for inferring co-accessibility, while simple binarization does not improve the predictions, as already suggested in (Martens *et al*., 2024). Additionally, while computing metacells can be interesting for reducing computational time, we observed a small drop in performance with both tested default strategies.

We also compared running time and memory usage (Figure 3.1d) for CIRCE and Cicero on both datasets, with and without computing metacells. On average, CIRCE ran almost 150 times faster and used 5.2 times less memory. On the largest dataset (PBMC 10X - single-cell data), CIRCE ran in 6 min 34 seconds instead of 1 day and 8 hours for Cicero, and used 35.8 GB of RAM instead of 223.1 GB.

Finally, to demonstrate CIRCE’s scalability and the new possibility it offers, we analysed a human atlas of fetal single-cell chromatin accessibility (Zhang *et al*., 2021). CIRCE processed the 720,616 cells and 1,041,455 DNA regions into a co-accessibility network in 42 min on an HPC. It identified 84,479,373 pairs of co-accessible regions. Since most of the computational time is actually spent in extracting small DNA windows from the initial input (columns of the AnnData input), we recommend using the csc_matrix format of scipy when analysing huge atlases since it is optimized for column extractions.

## 4. Conclusion

Here we present CIRCE, a Python package predicting cis-regulatory DNA interactions from single-cell chromatin accessibility data. CIRCE offers an optimised implementation of the algorithm proposed in (Pliner *et al*., 2018), adapted to the new challenges of recent single-cell datasets and compatible with the scverse environment. While providing very similar results to Cicero, CIRCE runs much faster and uses less memory, and can highly simplify the integration of cis-regulatory DNA interaction networks in Python workflows.

We also provide new insights on the preprocessing of single-cell ATAC and the use of metacells to infer co-accessibility with this algorithm. We reveal that binarization is unnecessary and that count normalization can be counter-productive. While different metacell strategies might actually improve the results by reducing noise, we didn’t observe such improvement from the tested strategies. However, they allow conserving good performances with a lower number of observations and reducing their sparsity. CIRCE’s optimized implementation will also permit analysing much bigger datasets without having to reduce the number of cells through aggregation. We envision that CIRCE will be a useful starting point for new gene activity inference tools in Python and integrated in the scverse ecosystem, which could leverage enhancer-promoter maps. Indeed, several methods propose considering chromatin accessibility of enhancers and promoters to estimate gene expression (Pliner et al., 2018).

## 5. Methods

### 5.1. Data preparation

#### BMMC NeurIPS dataset

The BMMC multiome used here was generated for the Open Problems challenge (Luecken *et al*., 2021) and is accessible under the GEO accession number <u>GSE194122</u>. We extracted the smallest batch of 1,771 cells and kept all the peaks expressed in at least one cell (110,235 peaks).

#### PBMC dataset

A PBMC multiome dataset was obtained from https://www.10xgenomics.com/datasets/pbmc-from-a-healthy-donor-granulocytes-removed-through-cell-sorting-10-k-1-standard-1-0-0. From the fragment files, we applied the same dataset preparation as in GRETA (Badia-i-Mompel *et al*., 2024). The peak calling and merging were achieved with Snapatac2 (Zhang *et al*., 2024), through the functions snap.tl.macs3, snap.tl.merge_peaks, and snap.pp.make_preaks_matrix. Only the cells expressing more than 100 genes in the scRNA-seq data were then kept. The final dataset contained 9,631 cells and 215,676 peaks.

#### PC-HiC dataset

The promoter capture HiC (PCHiC) dataset used to evaluate DNA region interactions prediction in the PBMC dataset is described in (Javierre *et al*., 2016). The processed interaction table is available as “PCHiC_peak_matrix_cutoff5.tsv” in the supplemental data S1: https://ars.els-cdn.com/content/image/1-s2.0-S0092867416313228-mmc4.zip.

#### Fetal human single-cell chromatin accessibility Atlas

The scATAC-seq atlas (Domcke *et al*., 2020), already preprocessed, was downloaded from https://scglue.readthedocs.io/en/latest/data.html under the name Domcke-2020.h5ad.

### 5.2. Reproducibility

The experiments are implemented as a Snakemake pipeline containing all the code to reproduce the experiments at https://github.com/cantinilab/circe_reproducibility. All the computations were realised on an HPC equipped with Linux Red Hat 8.8 and 2 AMD EPYC 7552 48-Core processors. For the benchmarking, CIRCE and Cicero were executed from their respective singularity containers (available in the reproducibility repository). Resource usage was limited to 20 cores and 430 GB of RAM through Snakemake rule configuration.

## 6. Acknowledgements

We acknowledge the help of the HPC Core Facility of the Institut Pasteur and Déborah Philipps for the administrative support.

## 7. Funding

The project leading to this manuscript has received funding from the European Union, ERC StG, MULTIview-CELL, 101115618 (L.C.). In addition, this work has been funded by the French government under the management of Agence Nationale de la Recherche as part of the “Investissements d’avenir” program, reference ANR-19-P3IA-0001; PRAIRIE 3IA Institute (L.C.) and by the Inception program “Investissement d’Avenir grant ANR-16-CONV-0005” (L.C.).

## Conflict of Interest

JSR reports in the last 3 years funding from GSK and Pfizer & fees/honoraria from Travere Therapeutics, Stadapharm, Astex, Owkin, Pfizer, Grunenthal, Tempus, and Moderna.

## Notes

### Summary of Updates

corrected error in names surnames order

## Bibliography

Badia-i-Mompel, P. et al. (2024) Comparison and evaluation of methods to infer gene regulatory networks from multimodal single-cell data. 2024.12.20.629764.

Badia-i-Mompel, P. et al. (2023) Gene regulatory network inference in the era of singlecell multi-omics. Nat. Rev. Genet.

Baran, Y. et al. (2019) MetaCell: analysis of single-cell RNA-seq data using K-nn graph partitions. Genome Biol., 20, 206.

Bilous, M. et al. (2024) Building and analyzing metacells in single-cell genomics data. Mol. Syst. Biol., 20, 744–766.

Blondel, V.D. et al. (2008) Fast unfolding of communities in large networks. J. Stat. Mech. Theory Exp., 2008, P10008.

Bravo González-Blas, C. et al. (2023) SCENIC+: single-cell multiomic inference of enhancers and gene regulatory networks. Nat. Methods, 20, 1355–1367.

Chari, T. and Pachter, L. (2023) The specious art of single-cell genomics. PLOS Comput. Biol., 19, e1011288.

Cusanovich, D.A. et al. (2018) A Single-Cell Atlas of In Vivo Mammalian Chromatin Accessibility. Cell, 174, 1309-1324.e18.

Dekker, J. et al. (2002) Capturing chromosome conformation. Science, 295, 1306–1311.

Domcke, S. et al. (2020) A human cell atlas of fetal chromatin accessibility. Science, 370, eaba7612.

Forrest, A.R.R. et al. (2014) A promoter-level mammalian expression atlas. Nature, 507, 462–470.

Fulco, C.P. et al. (2019) Activity-by-contact model of enhancer–promoter regulation from thousands of CRISPR perturbations. Nat. Genet., 51, 1664–1669.

Granja, J.M. et al. (2021) ArchR is a scalable software package for integrative single-cell chromatin accessibility analysis. Nat. Genet., 53, 403–411.

Grover, A. et al. (2024) UniversalEPI: Harnessing Attention Mechanisms to Decode Chromatin Interactions in Rare and Unexplored Cell Types. 2024.11.22.624813.

Hagberg, A.A. et al. (2008) Exploring Network Structure, Dynamics, and Function using NetworkX. Proc. 7th Python Sci. Conf. SciPy 2008, 11–15.

Hsieh, C.-J. et al. (2014) QUIC: Quadratic Approximation for Sparse Inverse Covariance Estimation. J. Mach. Learn. Res., 15, 2911–2947.

Javierre, B.M. et al. (2016) Lineage-Specific Genome Architecture Links Enhancers and Non-coding Disease Variants to Target Gene Promoters. Cell, 167, 1369-1384.e19.

Kamimoto, K. et al. (2023) Dissecting cell identity via network inference and in silico gene perturbation. Nature, 614, 742–751.

Kobak, D. and Linderman, G.C. (2021) Initialization is critical for preserving global data structure in both t-SNE and UMAP. Nat. Biotechnol., 39, 156–157.

Laska, J. and Narayan, M. (2017) skggm 0.2.7: A scikit-learn compatible package for Gaussian and related Graphical Models.

Lieberman-Aiden, E. et al. (2009) Comprehensive mapping of long-range interactions reveals folding principles of the human genome. Science, 326, 289–293.

Luecken, M. et al. (2021) A sandbox for prediction and integration of DNA, RNA, and proteins in single cells. Proc. Neural Inf. Process. Syst. Track Datasets Benchmarks, 1.

Martens, L.D. et al. (2024) Modeling fragment counts improves single-cell ATAC-seq analysis. Nat. Methods, 21, 28–31.

Pedregosa, F. et al. (2011) Scikit-learn: Machine Learning in Python. J. Mach. Learn. Res., 12, 2825–2830.

Persad, S. et al. (2023) SEACells infers transcriptional and epigenomic cellular states from single-cell genomics data. Nat. Biotechnol., 41, 1746–1757.

Pliner, H.A. et al. (2018) Cicero predicts cis-regulatory DNA interactions from single cell chromatin accessibility data. Mol. Cell, 71, 858-871.e8.

Sakaue, S. et al. (2024) Tissue-specific enhancer–gene maps from multimodal single-cell data identify causal disease alleles. Nat. Genet., 56, 615–626.

Sanborn, A.L. et al. (2015) Chromatin extrusion explains key features of loop and domain formation in wild-type and engineered genomes. Proc. Natl. Acad. Sci. U. S. A., 112, E6456–6465.

Sexton, T. et al. (2012) Three-dimensional folding and functional organization principles of the Drosophila genome. Cell, 148, 458–472.

Sheth, M.U. et al. (2024) Mapping enhancer-gene regulatory interactions from single-cell data. 2024.11.23.624931.

Stuart, T. et al. (2021) Single-cell chromatin state analysis with Signac. Nat. Methods, 18, 1333–1341.

Trimbour, R. et al. (2024) Molecular mechanisms reconstruction from single-cell multiomics data with HuMMuS. Bioinformatics, 40, btae143.

Virshup, I. et al. (2024) anndata: Access and store annotated data matrices. J. Open Source Softw., 9, 4371.

Virshup, I. et al. (2023) The scverse project provides a computational ecosystem for single-cell omics data analysis. Nat. Biotechnol., 41, 604–606.

Yao, Z. et al. (2021) A transcriptomic and epigenomic cell atlas of the mouse primary motor cortex. Nature, 598, 103–110.

Zhang, K. et al. (2024) A fast, scalable and versatile tool for analysis of single-cell omics data. Nat. Methods, 21, 217–227.

Zhang, K. et al. (2021) A single-cell atlas of chromatin accessibility in the human genome. Cell, 184, 5985-6001.e19.

Zhang, L. et al. (2022) DIRECT-NET: An efficient method to discover cis-regulatory elements and construct regulatory networks from single-cell multiomics data. Sci. Adv., 8, eabl7393.

